# Vectorized Capsid Rendering in the Browser with Capsid.js

**DOI:** 10.1101/2020.12.02.408252

**Authors:** Daniel Antonio Negrón

**Affiliations:** George Mason University, Bioinformatics and Computational Biology, 10900 University Blvd, Manassas, VA, 20110

## Abstract

**Motivation:** Few online services exist for rendering high-quality viral capsid figures compatible with common productivity software to develop effective infographics in the field of virology.

**Results:** The capsid.js library renders class I viral capsids within an online application that parameterizes style options, perspectives, and lattice patterns with SVG export.

**Availability:** This project is actively developed on GitHub (https://github.com/dnanto/capsid), distributed under the MIT License, hosted on GitHub Pages, and runs on modern browsers (https://dnanto.github.io/capsid/capsid.html).

**Supplementary information:** Supplementary data are available on GitHub.

## 1 Introduction

Molecular biology is the study of the building blocks of life and how they organize into complex objects via intricate processes and cycles. The field often condenses complicated concepts into simplified diagrams and cartoons for effective communication. Thus, such representations are useful for describing viral capsid structure and assembly.

The ViralZone web resource takes a direct approach, providing a static reference diagram for each known virus (Hulo *et al*., 2011). This contrasts with dynamic computational resources such as VIPER, which assembles realistic capsid structures from PDB files (Reddy *et al*., 2001). VIPERdb hosts the Icosahedral Server, but it only exports three-dimensional files in the same specialized format (Carrillo-Tripp *et al*., 2009). Neither resource provides an interactive online method for generating simple, two and three-dimensional structure files that are compatible with office software.

The capsid.js application is an interactive tool that renders icosahedral capsids and their nets in the browser. It parameterizes styling, projections, and lattice geometries and exports resulting models to SVG or PNG. The primary design goal is the generation of SVG images since they are infinitely scalable and compatible with word processors and vector graphics editors. Accordingly, this format is desirable for generating publication-quality figures. This means that the user is free to remix the SVG shape components.

## 2 Methods

### 2.1 Implementation

The application consists of an object-oriented JavaScript library that implements Caspar-Klug theory to generate and project the face of the unit icosahedron (Caspar and Klug, 1962). It also includes the Paper.js library for 2D vector graphics methods. A simple linear algebra engine provides methods to compute a camera matrix and project points.

### 2.2 Face

The first rendering step generates a grid of the selected tile geometry, including hexagonal, trihexagonal, snub hexagonal, rhombitrihexagonal, or the dual projection of any of the former. These additional tiles account for viruses with specialized lattice architectures (Twarock and Luque, 2019). Next, the procedure draws an equilateral triangle based on the h and k parameters, moving h tile units forward and k to the left or right depending on the selected levo or dextro parameter. This triangle serves as a cookie-cutter, removing the portion of the grid that serves as the face.

### 2.3 Color

The application includes color options for the hexamer, pentamer, fiber, and fiber knob domain. The procedure maps additional colors based on the selected tiling. Disambiguation between hexamer and pentamer determines whether the tile occurs within the it’s corresponding circumradius at each vertex of the enclosing face triangle.

### 2.4 Projection

To project each face, the procedure computes the unit icosahedron and applies the camera matrix to each vertex. Sorting each face by z-order achieves realistic occlusion. An affine transform operation then translates each face to the corresponding 2D projection coordinates of each face on the solid. To render vertex proteins, such as the adenovirus fiber, the procedure computes the vertexes of a larger icosahedron and includes the corresponding objects in the z-ordering operation.

**Fig. 1.**
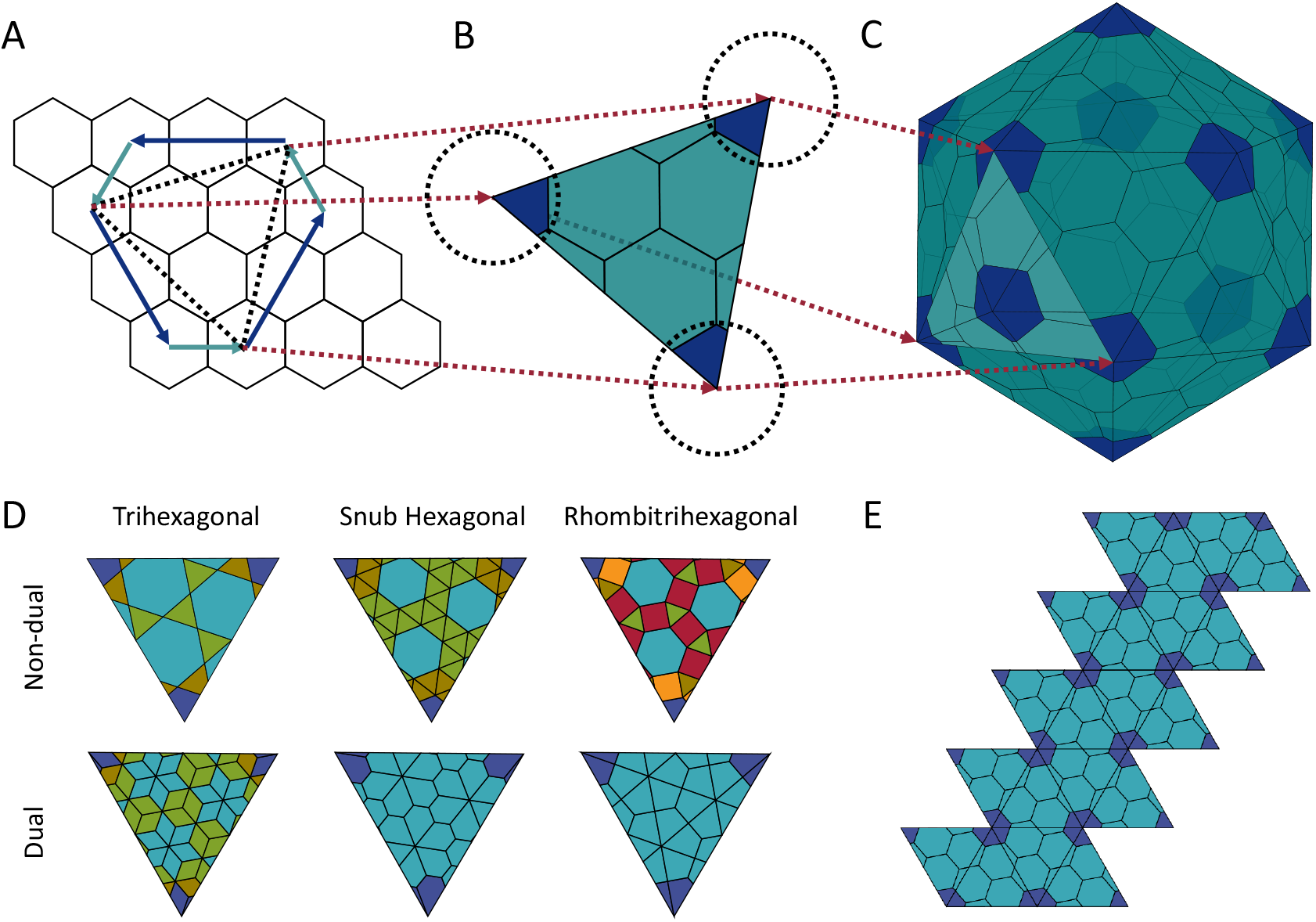
Construct the face and icosahedron for a levo T7 viral capsid (*T* = *h*^2^ + *hk* + *k*^2^ = 2^2^ + (2)(1) + 1^2^). This figure shows how office software such as Microsoft PowerPoint can import and convert the SVG files generated with capsid.js into shape objects. (A) Select the unit tile (hexagon) and draw a grid. Draw a triangle over the grid, moving h and k tile units forwards and left respectively. (B) Extract the triangular area from the grid. Color the hexamers and pentamers based on the unit tile circumradius from each face vertex. (C) Project the face to the 2D coordinates of the icosahedron based on the camera matrix. (D) Alternate tiling patterns. (E) The icosahedral net.

## 3 Discussion

The capsid.js app renders customizable icosahedral viral capsids. Development is active with plans to include prolate/oblate capsids. High-resolution SVG continues being a primary design goal to aid in the creation of detailed figures. This includes rendering all details as separate shapes. As a result, the current implementation may lag for large values of h or k. However, performance enhancements are possible by improving the rendering algorithm or switching to a GPU-based library, such as WebGL.

## Acknowledgements

The author would like the thank his dissertation committee: Dr. Donald Seto, Dr. Patrick Gillevet, and Dr. Sterling Thomas. Also, the author would like to thank Shane Mitchell, Mychal Ivancich, and Mitchell Holland for comments and review. This constitutes a portion of Chapter 2 of the PhD dissertation “Molecular Clock Analysis of Human Adenovirus” submitted to GMU.

